# Differential gene expression reveals host factors for viral shedding variation in mallards (*Anas platyrhynchos*) infected with low-pathogenic avian influenza virus

**DOI:** 10.1101/2021.08.23.457327

**Authors:** Amanda C. Dolinski, Jared J. Homola, Mark D. Jankowski, John D. Robinson, Jennifer C. Owen

## Abstract

Intraspecific variation in pathogen shedding impacts disease transmission dynamics; therefore, understanding the host factors associated with individual variation in pathogen shedding is key to controlling and preventing outbreaks. In this study, ileum and bursa of Fabricius tissues of mallards (*Anas platyrhynchos*) infected with low-pathogenic avian influenza (LPAIV) were evaluated at various post-infection time points to determine genetic host factors associated with intraspecific variation in viral shedding. By analyzing transcriptome sequencing data (RNA-seq), we found that LPAIV-infected mallards do not exhibit differential gene expression compared to uninfected birds, but that gene expression was associated with viral shedding quantity early in the infection. In both tissues, immune genes were mostly up-regulated in higher shedding birds and had significant positive relationships with viral shedding. In the ileum, host genes involved in viral cell entry were down-regulated in low shedders one day post-infection (DPI), and host genes promoting viral replication were up-regulated in high shedders on two DPI. Our findings indicate that viral shedding is a key factor for gene expression differences in LPAIV-infected mallards, and the genes identified in this study could be important for understanding the molecular mechanisms driving intraspecific variation in pathogen shedding.

## Introduction

Heterogeneity in pathogen transmission across individuals can impact the magnitude and duration of a disease outbreak (1, 2). Some individuals in a population are super-spreaders; that is, they are disproportionately responsible for secondary transmission cases of a contagious pathogen through increased contact rate and/or higher infectiousness (3). The super-spreader phenomenon can be traced back to Typhoid Mary in the early 1900s (4), humans infected with human immunodeficiency virus (HIV) in the 1980s (5, 6), wildlife spillover events in recent decades (4, 7), and more recently, in human outbreaks of novel coronaviruses (8). Super-spreading events are more often attributed to higher contact rates by some infected individuals in a population (2,9,10) rather than to variation in infectiousness; however, for pathogens that can be transmitted without direct contact from infected individuals, it stands to reason that pathogen shedding magnitude and duration may be as important as contact rate in generating super-spreading events. The individuals with the highest pathogen shedding, i.e. the “super-shedders”(11), have been shown to be drivers of super-spreading events as demonstrated in cattle infected with *Escherichia coli* O157 (12) and *Brucella abortus* (13), which are host-pathogen systems with indirect sources of transmission. Despite the apparent importance of individual variation in pathogen shedding to transmission dynamics, we know little about the mechanisms underlying differential pathogen shedding.

Identifying the extrinsic and intrinsic factors that affect an individual host’s pathogen shedding can improve our understanding of mechanisms generating and perpetuating super-spreading events. Extrinsic host factors linked to heterogeneity in pathogen shedding include food availability (14, 15), geographical location (16), toxicant exposure (17), and myriad environmental factors that can affect host immune competence through chronically elevated stress hormones (18). Intrinsic factors linked to pathogen shedding are a host’s age (19), sex (20, 21), physiology (22–24), and the immune response (24–26). In particular, observing how intraspecific variation in pathogen shedding relates to variation in host biological processes can improve our understanding of host factors involved in pathogen transmission.

The advancement of transcriptomics and gene expression studies have further highlighted the importance of the host immune response and physiology to variation in pathogen shedding. For instance, evaluating the bovine intestinal tract transcriptome revealed that T-cell responses and cholesterol metabolism were associated with super-shedding of *E. coli* O157 (24). Another study found that viral load in rainbow trout (*Oncorhynchus mykiss*) infected with infectious hematopoietic necrosis virus was correlated with type I and type II interferon transcript levels early in the infection (25). Analyzing gene expression simultaneously with pathogen shedding improves our understanding of infectious disease progression and the mechanisms driving transmission dynamics, but due to limited studies in only a few host-pathogen systems, further investigation is warranted.

Gene expression studies have also highlighted the value of examining multiple tissues at various time points post-infection since immune responses are tissue-specific and change over time (27–29). Successful viral infections, for example, are dependent upon cellular entry and the ability to evade the host immune response. The host provides initial protection by activating the innate immune response through pattern recognition receptors (PRR) that activate the transcription of interferons, which leads to the production of antiviral interferon-stimulated genes (ISGs) (30, 31). This process provides protection to the host since many ISGs are known to inhibit virus cell entry and replication, degrade viral RNA, regulate cell apoptosis, and have regulatory effects on the interferon pathway (31, 32). The host also stimulates immune cell dependent mechanisms (cellular immunity) to eliminate infected cells and activate adaptive immunity consisting of antigen recognition by major histocompatibility complex (MHC) molecules, T-cell activation, and B-cell activation for the production of antibodies later in the infection (33–35). Depending on the type of tissue infected, the innate and adaptive immune systems respond differently (29), thus it’s important to evaluate tissue-specific host gene expression at various time-points post-infection.

In this study, we analyzed the association between gene expression and viral shedding variation in avian influenza virus (AIV) infected mallards (*Anas platyrhynchos*). AIVs are type A influenza viruses that can spillover into domestic swine and poultry, thus AIVs pose a significant health risk to domestic animal and human populations (36). Wild waterfowl are the natural reservoirs for low-pathogenic avian influenza virus (LPAIV), a virus transmitted via the fecal-oral route directly or indirectly through AIV contaminated water sources (37). Previous studies have shown that LPAIV-infected mallards exhibit extreme heterogeneity in pathogen shedding (38), which has been linked to both extrinsic (food availability) (15) and intrinsic (gut physiology) (22) host factors. Yet, these factors alone do not explain all the observed variation in viral shedding. Previous gene expression studies of AIV-infected poultry consistently show that gene expression is up-regulated in AIV-infected birds, many of which are associated with the immune response (28,39–43). To our knowledge, however, a longitudinal transcriptome-wide study is yet to be conducted to analyze the association between gene expression and viral shedding.

We hypothesized that gene expression in the ileum and bursa would be associated with intraspecific viral shedding variation in LPAIV-infected mallards because LPAIV primarily replicates in the epithelial cells of both the intestines and bursa of Fabricius (hereafter referred to as “bursa”) of waterfowl by gaining entry via sialic acid receptors (44–47). Further, the antiviral immune response occurs at the site of replication (31,48,49); therefore, we predicted that gene expression would differ: 1) between infected and uninfected birds, 2) between days post-infection, with innate immunity genes expressed early in the infection and adaptive immunity genes expressed later in the infection, 3) between bird groups with various viral shedding levels, and 4) that viral shedding magnitude would be correlated with the expression level of genes involved in cell entry and the immune response. We tested our hypotheses by evaluating gene expression at various time points post-infection to detect differential gene expression with respect to infection stage, infection site, and viral shedding in an important reservoir host for AIV, the mallard (50–53). In this study we further our understanding about the genes and pathways associated with individual heterogeneity in pathogen shedding.

## Materials and methods

### Ethics statement

Eggs were collected under the U.S. Fish and Wildlife Permit (M Bl 94270-2) and North Dakota Game and Fish Department License #GNF03639403. All capture, handling, infections, and sampling of mallards was approved by the Michigan State University (MSU) Institutional Animal Care and Use Committee (AUF 12/16-211-00). All euthanasia procedures, including the use of pentobarbital sodium and phenytoin sodium solution (Beuthanasia-D Special, Merck Animal Health, Madison, NJ, USA) and CO_2_ inhalation, were in accordance with the Animal Welfare Act and Guidelines to the Use of Wild Birds in Research (54).

### Birds and virus

Wild mallards collected *in ovo* were used for this study. Analyzing the gene expression of wild birds is important due to the inherent genetic variation within the population (55), a likely factor driving variation in a species response to viral infection (15). Eggs were collected from hen nests located in uncultivated fields of Towner County, North Dakota, USA (48.44, -99.31) in May - June 2015, and shipped to MSU where they were incubated (Sportsman 1502 Egg Incubator, GQF Manufacturing Co., Savannah, GA) at 37.5°C with 45-50% humidity until hatched. Full details of egg collection and raising ducks can be found in Dolinski et al. (22).

LPAIV A/northern pintail/California/44221-761/2006 (H5N9) was acquired from the USGS National Wildlife Health Center (NWHC) in Madison, WI (USDA Veterinary Permit 44372). The virus was propagated in embryonated pathogen free eggs (Charles River, Norwich, CT, USA) under biosafety level two laboratory conditions at MSU (56). The propagated virus (stock) was sent to USGS NWHC and determined to have a viral titer of 7.63 log EID_50_/ml using the Reed-Munch 50% egg infectious dose method (57).

### Experimental design

Once all the birds were eight weeks or older, they were moved to individual cages (76×61×41cm) and randomly assigned to one of seven experimental groups using a random-pseudostratified method controlling for age, weight, and nest. Group identification and size are shown in Fig 1, with the group name referring to infection status and the day post-infection (DPI) when the group was sacrificed for tissue sample collection.

**Fig 1.**
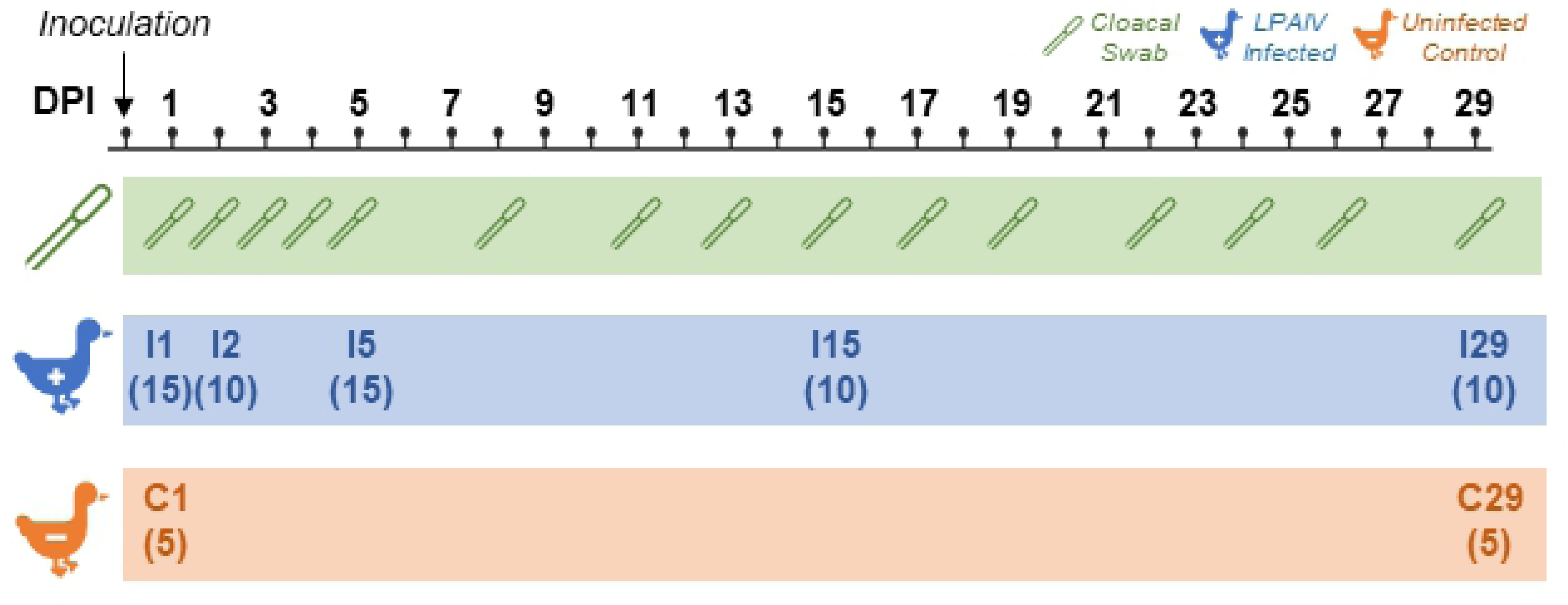
Experimental timeline of sample collection days post-infection (DPI). Groups are designated by infection status (I = LPAIV-infected, C = uninfected control), DPI sacrificed and (N) quantity of birds in each group. Ileum and bursa were collected upon sacrifice. Cloacal swabs were collected from all living birds designated with the swab icon.

Mallards (n = 60) were inoculated with 5.63 log EID_50_/ml of LPAIV H5N9 in Dulbecco’s Modified Eagle Medium (DMEM; Gibco® by Life Technologies, Grand Island, NY, USA) and the remaining mallards (n = 10) were sham inoculated with sterile DMEM. Inoculum (1.0 mL) was administered to birds with one drop in each eye and nare, then the rest dispensed in the bird’s esophagus.

Cloacal swabs were used for quantification of virus shedding (virus titers), because virus in the cloaca represents transmissible virus and the sample can be repeated over time (58). Cloacal swabs were collected on days shown in Fig 1 using cotton-tipped swabs (Puritan, Guilford, ME), placed in 3.0 mL of brain-heart infusion broth (Sigma-Aldrich, St. Louis, MO, USA), and immediately placed on wet ice and stored at -80°C within three hours after collection.

Birds in each group were euthanized and necropsied at intervals illustrated in Fig 1. Birds sacrificed on one DPI (I1, C1) were euthanized via lethal injection with pentobarbital sodium and phenytoin sodium solution (Beuthanasia-D Special, Merck Animal Health, Madison, NJ, USA), but due to complications with intravenous injections, all other birds were sacrificed by CO_2_ inhalation. Birds were stored on beds of ice until necropsies were conducted one to six hours after being euthanized. During necropsies, tissue sections (2 mm^3^) of ileum and bursa were collected, and the coelomic cavity was assessed for gross pathology. Each tissue section was placed in 1.0 mL of room temperature RNA stabilizing solution (RNAlater®, Sigma-Aldrich, St. Louis, MO, USA), then removed from the solution after 24 hours and stored at -80°C.

### Viral RNA extraction and quantification from cloacal swabs

Cloacal swab samples were thawed at room temperature and viral RNA was extracted from 200 μL of each sample using the MagMAX™-96 AI/ND Viral RNA Isolation Kit (Applied Biosystems® by Thermo Fisher Scientific, Vilnius, Lithuania) with modifications (59).

Extracted viral RNA was stored in 500 μL of elution buffer. The virus titer for each cloacal swab sample was quantified by real time reverse transcription-quantitative polymerase chain reaction (qPCR) targeting the matrix protein gene. We used the TaqMan® RNA-to-Ct™ 1-Step Kit (Applied Biosystems® by Thermo Fisher Scientific, Foster City, CA, USA), primers/probe specified by Spackman et. al (60), and 2 μL of sample RNA in three replicates. CT values were used to estimate virus quantity as EID_50_/mL via standard curve (QuantStudio™ 7 Flex Real-Time PCR Software System v1.3) of stock viral RNA (7.63 log EID_50_/ml) in a series of 10-fold dilutions. The three replicates per sample were averaged to derive a single estimated virus titer for each sample.

### Determining shed level groups

Previous analysis of virus titer data in mallards determined that 99% of cumulative LPAIV was detected during the first five DPI (22), and birds were observed to disproportionately shed LPAIV according to the 20/80 rule (38); therefore, birds in LPAIV-infected groups I1, I2, and I5 were subdivided into three shed level groups (Fig 2). High shedders (H) were the top 20% of birds with the highest cloacal virus titer, moderate shedders (M) were the top half of the lowest 80% of virus titers, and low shedders (L) were the bottom half of the lowest 80% of virus titers in each group. To detect differential gene expression related to active viral shedding during the innate immune response (48, 49), only the virus titer collected on the day the bird was sacrificed was used to categorize shed level groups in early-stage infection (groups I1 and I2). To detect differential gene expression related to the adaptive immune response (35, 61), cloacal virus titers on one to five DPI were averaged to create a cumulative virus titer for late-stage infection (group I5).

**Fig 2.**
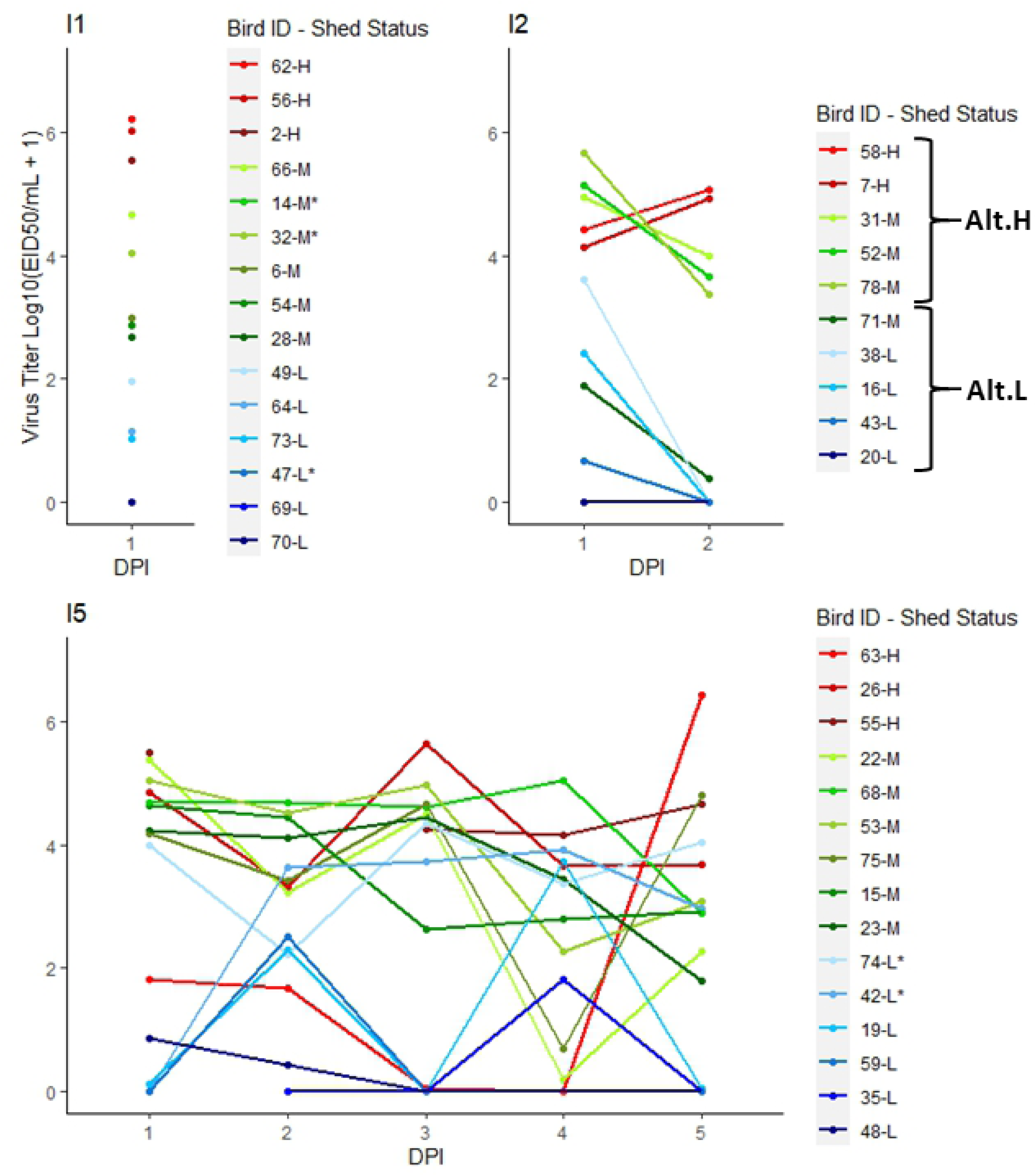
Cloacal swab virus titer profiles for LPAIV-infected mallards sacrificed on one (I1), two (I2), and five (I5) days post-infection (DPI). Shed status of high (H), moderate (M), or low (L) was assigned to individuals by their last virus titer (I1, I2) or the average virus titer across all DPI (I5). Alternative high (Alt.H) and alternative low (Alt.L) shed status groupings were assigned to individuals in I2 based on cumulative virus titers clustered across one and two DPI. Due to low host RNA quality, *individuals were not included in ileum differential gene expression analyses.

After evaluating virus titer data, we observed that group I2 cumulative virus titer data clustered into two distinct shed level groups (Fig 2); therefore, we also conducted differential gene expression analysis using an alternative two-group shed level classification of high shedders (Alt.H) and low shedders (Alt.L) for group I2.

### Tissue mRNA extraction, cDNA library preparation, and sequencing

Total mRNA was extracted from ileum and bursa tissue (15-30 mg) using the Qiagen RNeasy mini kit (QIAGEN®, Hilden, Germany) following the manufacturer’s protocol. Total mRNA was stored in 500 μL of RNase free water at -80°C.

RNA quality was determined prior to library preparation using a Qubit® Fluorometer 1.0 (Molecular Probes, Life Technologies, Eugene, OR, USA). Sixty-six of 70 bursa samples had RNA integrity number (RIN) scores >8.0 and one bursa sample had a RIN score of <7.0 (S1 Table). All 70 bursa samples were sequenced at the MSU Research Technology Support Facility. Thirty of the 70 ileum samples had RIN scores >8.0 and 35 had RIN scores <7.0. Library preparation and RNA sequencing was performed at MSU for 22 ileum samples with RIN scores >8.0 in groups I2, I5, and C1. Due to lack of availability at MSU, library preparation and RNA sequencing for an additional 20 ileum samples with RIN scores between 4.9 and 9.4 was conducted at the University of Minnesota (UMN) Genomics Center in St. Paul, MN.

Sequencing libraries were prepared using the NuGEN Ovation® Universal RNA preparation kit (Tecan Genomics, Chesapeake Drive, CA, USA) according to the manufacturer’s specifications. Custom probes complementary to mallard rRNA sequences were designed and used for the Insert Dependent Adaptor Cleavage (InDA-C) portion of the protocol. Completed library quality was assessed using Qubit® dsDNA HS (Molecular Probes, Life Technologies, Eugene, OR, USA) and Caliper LabChip® GX DNA HS (Caliper, A PerkinElmer Company, Hopkinton, MA, USA) assays. Libraries sequenced at MSU were combined into six pools for multiplexed sequencing, including five pools of 16 libraries each, and a sixth pool of 13 libraries. Library pools were quantified using Kapa Biosystems Illumina Library qPCR assay prior to sequencing. Each pool was loaded on one lane of an Illumina HiSeq 2500 High Output flow cell (v4) and sequencing was performed in a 2×150bp paired end format using HiSeq SBS reagents. Libraries at UMN were combined into a single pool and sequenced in one lane of an Illumina NovaSeq S1 2×150-bp run. Base calling was done by Illumina Real Time Analysis (RTA) v1.18.64 and output of RTA was demultiplexed and converted to FastQ format with Illumina Bcl2fastq v1.8.4.

### RNA-seq read processing

Quality of raw cDNA reads was assessed using FastQC (62) prior to trimming and filtering. Sequence regions with a quality score below 15 and sequences shorter than 40bp were removed using Trimmomatic (63). FastQC was used again to verify quality and trimming.

Sequences were aligned to the most recent mallard reference genome, Ensembl ASM874695v1 (64), using STAR (65, 66). Gene-specific read counts for bursa samples were acquired with HT-seq (67) using the STAR output, then RSEM (68) was used to estimate read abundance. Due to low RIN scores of 14 ileum samples (RIN = 4.9 – 7.3) likely caused by time delays between euthanasia and necropsy (69), we used DegNorm (70) to account for effects of degradation in ileum samples. Low RIN scores (RIN <8.0) can negatively impact analyses by misaligning sequences resulting in inaccurate transcript counts (71, 72); however, DegNorm adjusts read counts for heterogeneity in mRNA sequence degradation on an individual gene basis while also accounting for sequencing depth (70).

### Differential expression analyses

We performed differential gene expression analyses using the R packages “EdgeR” (73, 74) and “limma” (75) in R version 4.0.0, using the approach outlined by Law et al. (76). Differential expression analysis was performed separately for each tissue and included gene- and transcript-level analyses for the bursa and only gene-level for the ileum, because isoform counts are not estimated by DegNorm.

Lowly expressed genes and transcripts were filtered by requiring expression of >0.5 counts per million in at least 25% of the birds. A false discovery rate (FDR) corrected alpha value of 0.1 and a log fold count difference (LFC) of 0.5 were required to establish differential expression. These thresholds were established to allow for detection of subtle differential gene expression as previously observed between LPAIV-infected and uninfected ducks (41–43). To account for sex-based differences and variation in sequencing pool, sex and pool were included as covariates in each analysis. Gene names (HGNC symbols) were assigned based on ENSEMBL annotation information that accompanied the reference genome. KEGG pathway analyses were performed using the “kegga” function of “limma”. Pathways with p-values <0.05 were determined as over-represented (enriched) pathways of differentially expressed genes or transcripts (DEG, DET) per analysis.

Preliminary differential expression analysis showed that ileum and bursa samples clustered based on tissue-specific gene expression (Section 1 in S2 Document). Improper tissue-based clustering for two individuals (1 and 72) in group C1 suggested improper sample labeling may have occurred in those cases; therefore, these two individuals were removed from analyses, leaving a total of 68 bursa samples and 40 ileum samples. Additionally, due to an absence of differential gene expression between C1 and C29 (Section 2 in S2 Document), we combined the two control groups into one group “C” for subsequent analyses. Sample size for each group in each differential expression analysis is provided in Table 1.

**Table 1.** Group sample size for differential gene expression analyses.

| Comparison | Tissue | Group |  |  |  |  |  |
| --- | --- | --- | --- | --- | --- | --- | --- |
|  |  | C | I1 | I2 | I5 | I15 | I29 |
| Infected vs control | Bursa | 8 | 15 | 10 | 15 | 15 | 10 |
|  | Ileum | 5 | 12 | 10 | 13 |  |  |
| Infection by DPI | Bursa |  | 15 | 10 | 15 | 15 | 10 |
|  | Ileum |  | 12 | 10 | 13 |  |  |
|  |  | Sample size per viral shedding group<br>High, Moderate, Low |  |  |  |  |  |
| Shed Level | Bursa |  | 3,6,6 | 2,4,4 | 3,6,6 |  |  |
|  | Ileum |  | 3,4,5 | 2,4,4 | 3,4,6 |  |  |

To address our hypothesis and predictions, differential gene expression comparisons were conducted separately for ileum and bursa tissues. For our prediction that infection status induces differential gene expression, we compared LPAIV-infected groups on each DPI to the uninfected control group. For our prediction that LPAIV infection induces differential gene expression over time, we also compared each LPAIV-infected group to each other. For our prediction that virus shed level induces differential gene expression, we conducted analyses for early (groups I1 and I2) and late-stage infection (group I5) by comparing high, moderate, and low shed level groups for each DPI. Additionally, an alternative analysis was conducted for group I2 using the Alt.L and Alt.H comparison.

Once analyses were complete, a literature search was conducted for DET/DEGs using gene names and search terms including “immune response,” “influenza virus,” and “viral infection” on google scholar. The search yielded a total of 65 peer-reviewed papers. A total of 37 peer-reviewed journal articles were associated with 36 DEGs involved with the immune response (77–79), and a total of 27 journal articles were associated with 39 DEGS that are host cell factors of influenza A virus replication (80–83), similar to influenza host factors (orthologs or paralogs), or were involved in the replication of other viruses.

### Candidate gene analysis

A candidate gene analysis was conducted to observe linear relationships between gene expression and LPAIV virus titers. We selected candidate genes based on what is broadly known about the host’s immune response to viral infections and more specifically about the avian response to AIV infection (31,39,42,84–87). We also selected genes associated with production of sialic acid receptors (i.e. sialyltransferase genes) (88), since sialic acid receptors bind to influenza A viruses and allow for cell entry (46). Seventy-six search terms were identified based on gene function or name (e.g., interferon, IFIT; Table A in S3 File) for an automated search of the ENSEMBL *Anas platyrhynchos* genome using ‘hgnc_symbol’ and ‘description’ fields. This search identified a list of candidate genes and transcripts that were manually filtered to only include relevant genes. Using only transcripts and genes that showed variability in expression, we used linear mixed models to assess the relationship between gene expression and a bird’s virus titer on the day they were sacrificed. Virus titers were adjusted to log_10_(virus titer +1) transformed values and gene expression was analyzed as log_2_ transformed. Each targeted gene and transcript were assessed separately using automated sub-setting and looping through the dataset. Statistical significance was assessed based on a 0.05 false discovery rate. Sex and DPI were included as fixed effects to account for sex and time-point related differences in gene expression, and sequencing pool was a random effect. P-values and conditional R^2^ were reported for each significant model, as well as each LPAIV-infected bird group. The “nlme” R package was used for all linear mixed model analyses (89).

## Results and discussion

All LPAIV-infected birds tested positive for LPAIV as demonstrated by at least one cloacal swab, ileum tissue, or bursa tissue sample with a Ct value <40 on qRT-PCR (22, 90), whereas all uninfected control birds tested negative. No individuals died prior to scheduled euthanasia or exhibited signs of disease, such as ruffled feathers, lethargy, or gross pathology, consistent with previous findings (91).

### RNA sequencing

Library preparations generated ≥750 M reads for each lane. Mean quality scores for all libraries were ≥Q30. Bursa tissue samples yielded an average of 9,261,147 raw reads (range: 4,141,232 – 14,172,345) and ileum samples yielded an average of 10,881,788 raw reads (range 6,737,825 – 17,963,033). After filtering, we retained an average of 8,984,110 bursa reads (range: 4,062,211 – 13,742,163) and an average of 10,526,592 ileum reads (range: 6,567,571 – 17,444,644) per individual for analyses.

### Differential gene expression between LPAIV-infected and uninfected control groups

Differential gene expression was not observed between LPAIV-infected and uninfected control birds at each DPI in the bursa or ileum (Section 3 in S2 Document), despite our lenient DEG screening criteria. In previous studies, differential gene expression was observed between AIV-infected and uninfected poultry (28,41–43,92); yet, in studies that looked at both LPAIV and highly pathogenic AIV (HPAIV), gene expression was greater in HPAIV-infected birds than LPAIV-infected birds (41–43). Unlike other gene expression studies that used captive/hatchery-bred mallards (28,41–43,92), we used wild-bred mallards which have been shown to exhibit greater LPAIV viral shedding variation compared to captive-bred mallards (15). The high interindividual viral shedding variation of wild-bred LPAIV-infected mallards may therefore have reduced our ability to detect a statistical difference in gene expression between infected and uninfected mallards. Interindividual viral shedding variation was not addressed in any of the previous AIV gene expression studies (28,41–43,92); therefore, it is not possible to state whether the variation in viral shedding reported here differs from other studies; however, previous genetic studies show that there are clear genetic differences between wild- and captive-bred ducks (93), including differences in digestive system morphology (94, 95). Given the genetic and viral shedding differences between wild-bred and captive-bred ducks, we recommend that future studies use wild birds to better represent transmission dynamics in nature.

### Differential gene expression of LPAIV-infected birds over time

Within LPAIV-infected bird groups only, we observed few DETs and DEGs over the course of the infection. In the bursa, one DET (*HSPA8*) and two DEGs (*FOSB*, *HSPA8*) were down-regulated in group I2 compared to group I29 (Sections 3B and 3C in S2 Document). *HSPA8* and *FOSB* had a similar expression pattern that was lowest on two DPI and gradually increased at each DPI thereafter (Fig 3). *FOSB* codes for a transcription factor subunit for AP-1 which regulates the expression of *IFN-γ* (96). *HSPA8* (aka hsc70) mediates viral production by binding to viral protein M1 and NP transporting these viral proteins from the nucleus to the cytoplasm after replication (97, 98). Only *HSPA8* was associated with four enriched KEGG pathways in spliceosome, MAPK signaling, protein processing in the endoplasmic reticulum, and endocytosis functions. Since the highest viral shedding quantities are observed on one, two, and three DPI (22), the expression profile of these two genes suggests that mallards may down-regulate *FOSB* and *HSPA8* in the bursa as a protective mechanism in response to LPAIV infection.

**Fig 3.**
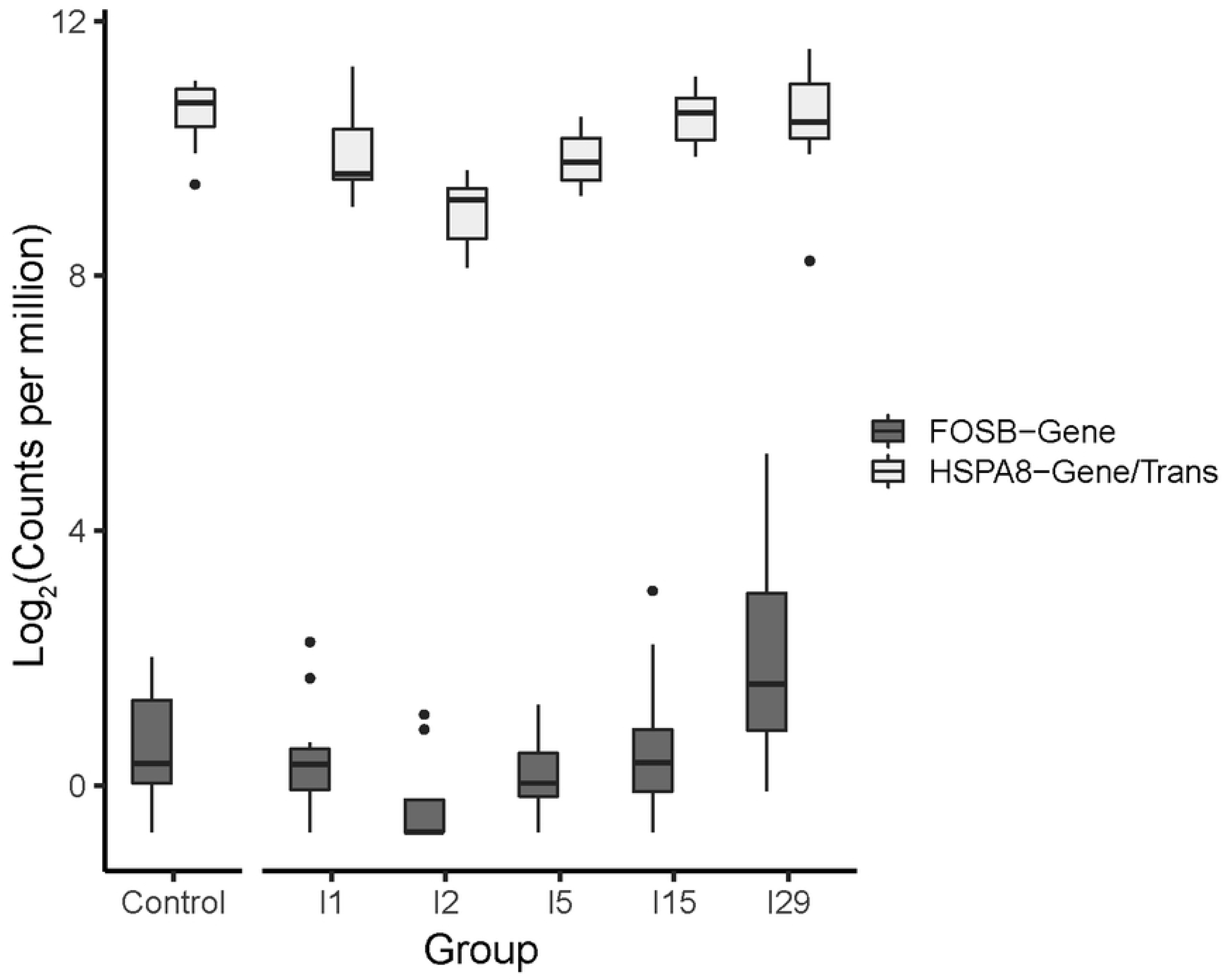
Expression of *FOSB* gene and *HSPA8* gene and transcript (“Gene/Trans”) in the bursa of uninfected control and LPAIV-infected groups. Infected groups I1, I2, I5, I15, and I29 correspond to the DPI birds were sacrificed. Differential expression (FDR <0.1, LFC >0.5) was only observed between I2 and I29. *HSPA8* had identical gene expression at the transcript (trans) and gene levels.

In the ileum, only one gene (*S100A12*) that codes for a calcium binding protein was differentially expressed and had lower expression in group I1 compared to group I2 (Section 3A in S2 Document). No KEGG pathways were enriched in the ileum.

### Differential gene expression between virus shed level groups during early stage infection

We observed differential gene expression between shed level groups early in the infection, with the majority of DEGs observed in the ileum. Between shed level groups in the ileum, we observed 406 DEGs in group I1 (Fig 4, Section 4A in S2 Document) and 186 DEGs in group I2 (Fig 4, Section 5A in S2 Document). In group I1, DEGs were only observed between low and moderate shedders with 97% of the DEGs up-regulated in the moderate shedders. In the ileum of group I2, DEGs were observed between high shedders and low/moderate shedders with 98% of the DEGs up-regulated in high shedders. In the alternate shed level comparison (Alt.L vs Alt.H) we observed 266 DEGs in group I2 (Fig 4, Section 6A in S2 Document) with 54% of the DEGs up-regulated in high shedders.

**Fig 4.**
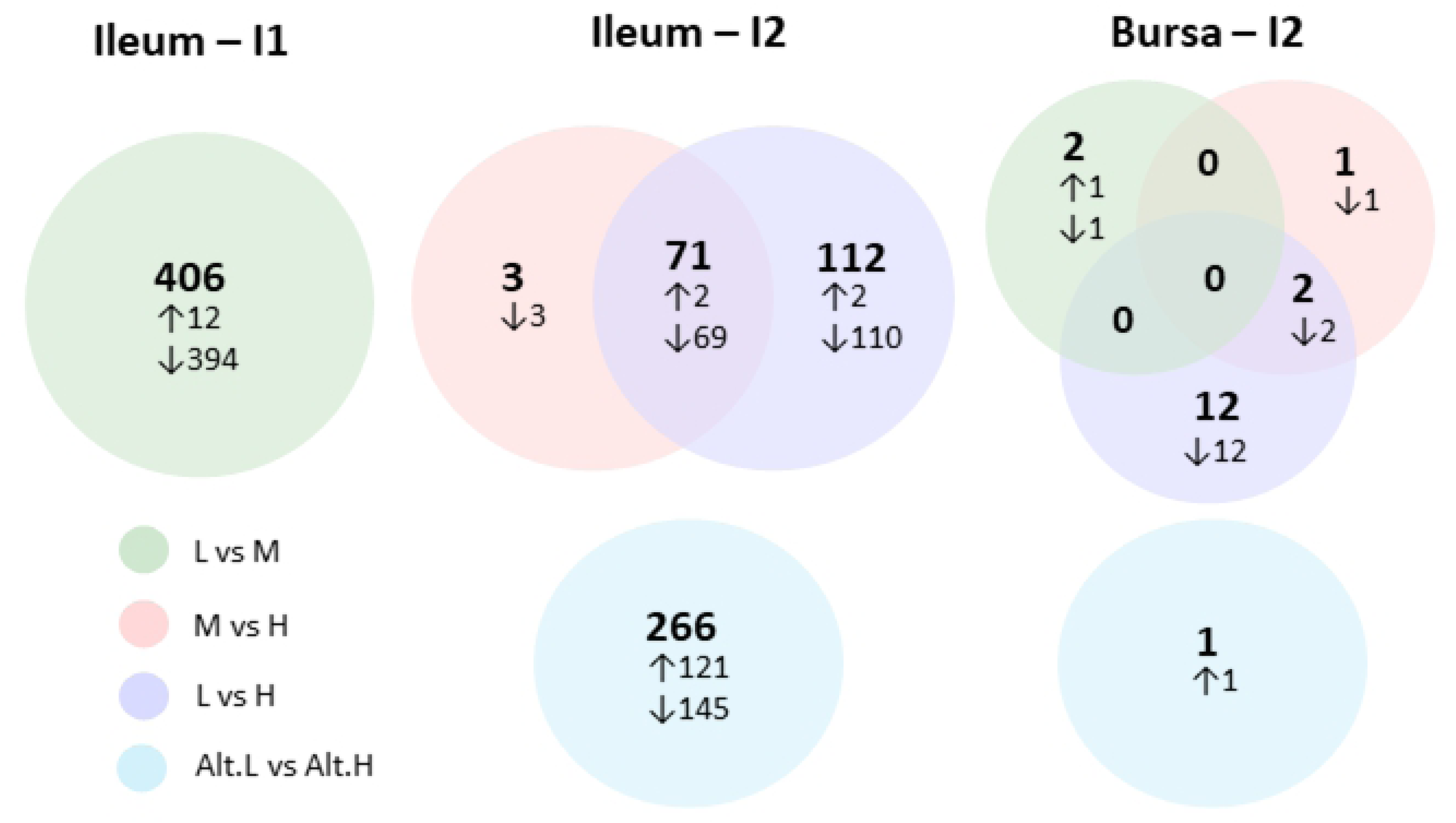
Number of differentially expressed genes (DEG) between shed level groups (low (L), moderate (M), high (H)) of LPAIV-infected mallards in the ileum and bursa on one (I1) and two (I2) DPI. An additional analysis for I2 divided birds evenly into alternative low and high shedders (Alt.L vs. Alt.H). Arrows reflect the number of DEGs that are up (↑) or down (↓) regulated in the first shed level group listed in each comparison.

In the bursa, we did not observe any DETs in group I1 (Section 4C in S2 Document) or group I2 (Section 5C and 6C in S2 Document). While no DEGs were observed between shed level groups in the bursa of group I1 (Section 4B in S2 Document), 17 DEGs were observed in group I2 (Fig 4, Section 5B in S2 Document), 88% of which were up-regulated in high shedders, a pattern consistent with DEGs in the ileum. In the bursa of the Alt.L vs. Alt.H group I2 comparison, we observed only one up-regulated gene in low shedders compared to high shedders (Fig 4, Section 6B in S2 Document). Fold count differences for all DEGs can be viewed in S2 Document.

KEGG pathway information was only available for DEGs between shed level groups in the ileum. In the ileum of group I1, KEGG pathways could be assigned to 51 out of the 406 DEGs observed which represented more than 50% metabolic pathways (Fig 5, Section 4A in S2 Document). Out of all group I1 KEGG pathways assigned, 11 metabolic and two cellular process pathways were enriched (p < 0.05), signifying likely biological significance of these pathways.

**Fig 5.**
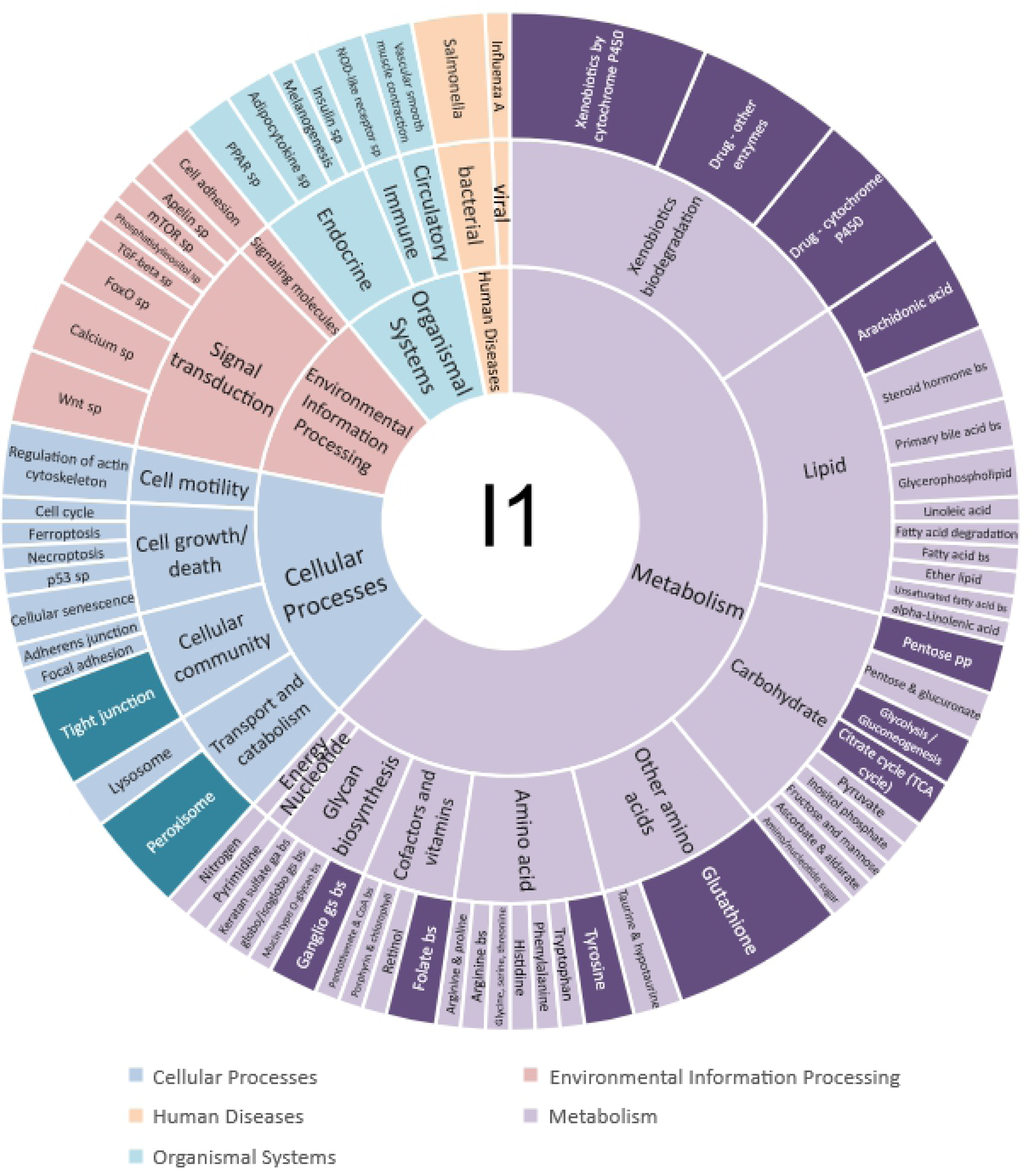
KEGG pathways of DEGs in the ileum between shed level group comparisons on one day post-infection (I1). KEGG pathways in darker colors indicate statistically enriched pathways (p < 0.05). Abbreviations: biosynthesis (bs), signaling pathway (sp), glycosaminoglycan (ga), glycosphingolipid (gs), phosphate pathway (pp).

Metabolic functions are commonly linked to viral infections (99), with lipid metabolism (100) and cellular kinases (101) most associated with influenza infections. In chickens infected with HPAIV H5N1, lipid metabolism was enriched on one DPI (41). It is suggested that lipid metabolism is important during the early stages of influenza infection because viral cell entry relies on membrane trafficking, and the cell membrane is made primarily of phospholipids (41). Interestingly, cholesterol metabolism was recently associated with super-shedding cattle infected with *E.coli* O157 (24), which is also an intestinal pathogen. These findings suggest that these metabolic pathways may be important host factors responsible for intraspecific variation of intestinal pathogens.

In the ileum of group I2, KEGG pathways could be assigned to 28 out of the 186 DEGs between the three shed level groups (Fig 6, Section 5A in S2 Document) and 31 out of the 266 DEGs between Alt.L and Alt.H (Fig 7, Section 6A in S2 Document). More than 50% of the enriched pathways in group I2 were involved in genetic information processing, which was not unexpected given that the influenza A viral replication cycle utilizes host cell machinery to replicate (102).

**Fig 6.**
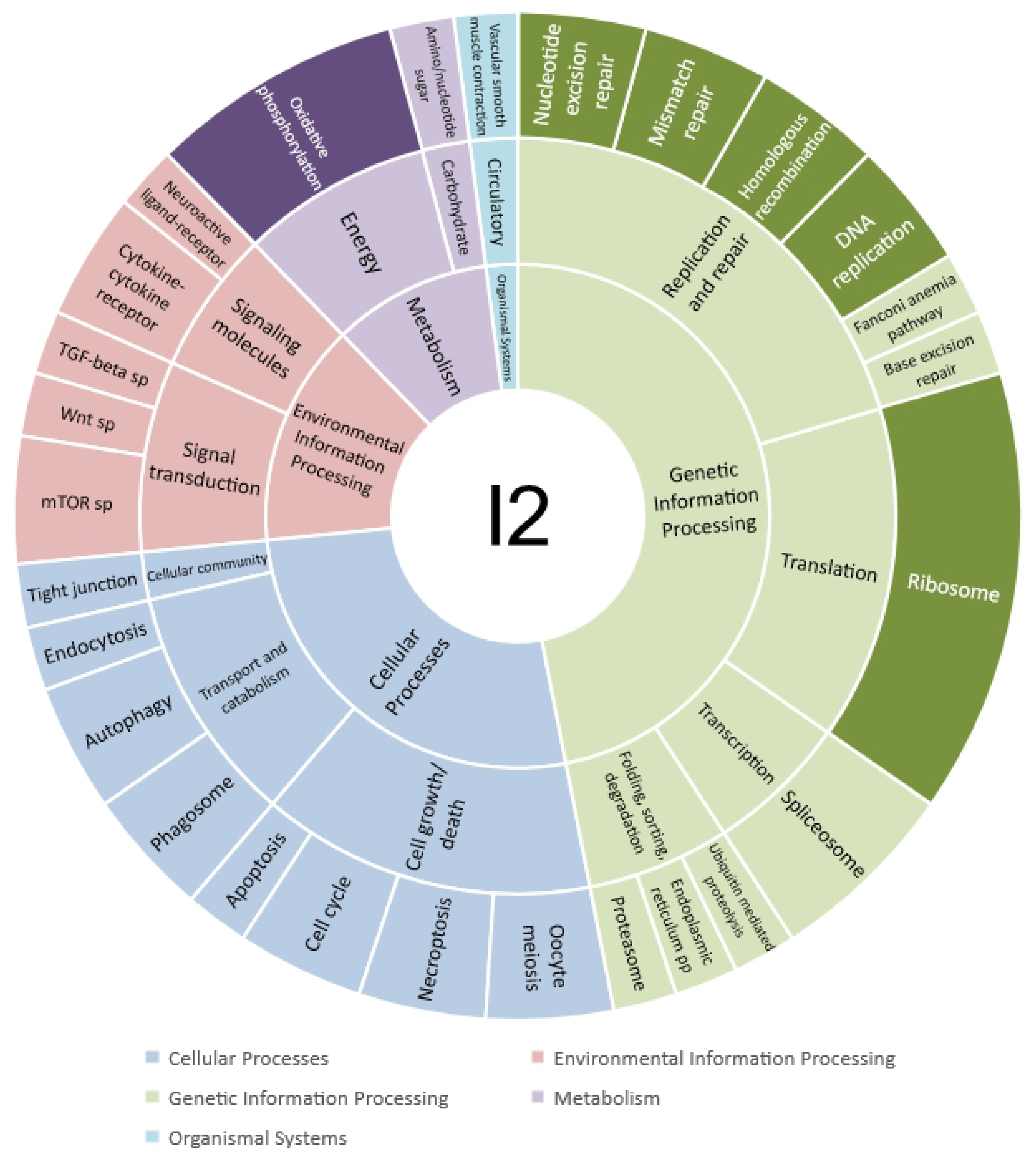
KEGG pathways of DEGs in the ileum between shed level group comparisons on two days post-infection (I2). KEGG pathways in darker colors indicate statistically enriched pathways (p < 0.05). Abbreviations: signaling pathway (sp), protein processing (pp).

**Fig 7.**
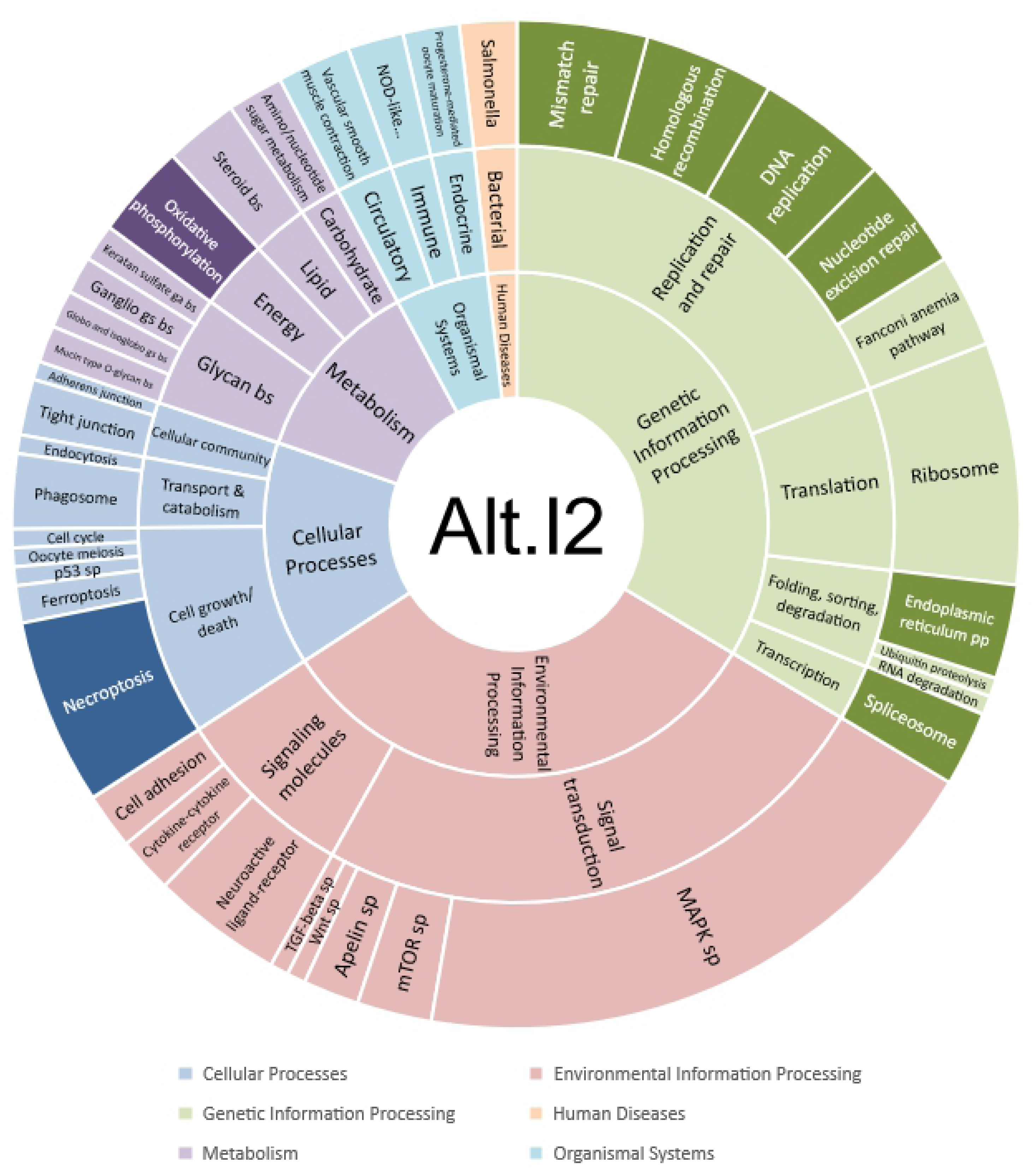
KEGG pathways of DEGs in the ileum between alternative shed level group comparisons (Alt.L vs. Alt.H) on two days post-infection (I2). KEGG pathways in darker colors indicate statistically enriched pathways (p < 0.05). Abbreviations: biosynthesis (bs), protein processing (pp), signaling pathway (sp), glycosaminoglycan (ga), glycosphingolipid (gs).

Genes of the immune system were differentially expressed between shed level groups early in the infection in both the ileum and the bursa. In the ileum of groups I1 and I2, genes with interferon pathway, cellular immunity, and apoptosis functions were up-regulated in higher shedding birds for 43 out of the 47 significant comparisons (Fig 8, Table 2). In the bursa of group I2, four interferon-stimulated genes were up-regulated in high shedders compared to low shedders (Table 3). The up-regulation of immune genes in higher shedding birds of I1 and I2 groups supports our prediction that viral shedding is closely associated with innate immunity early in the infection.

**Fig 8.**
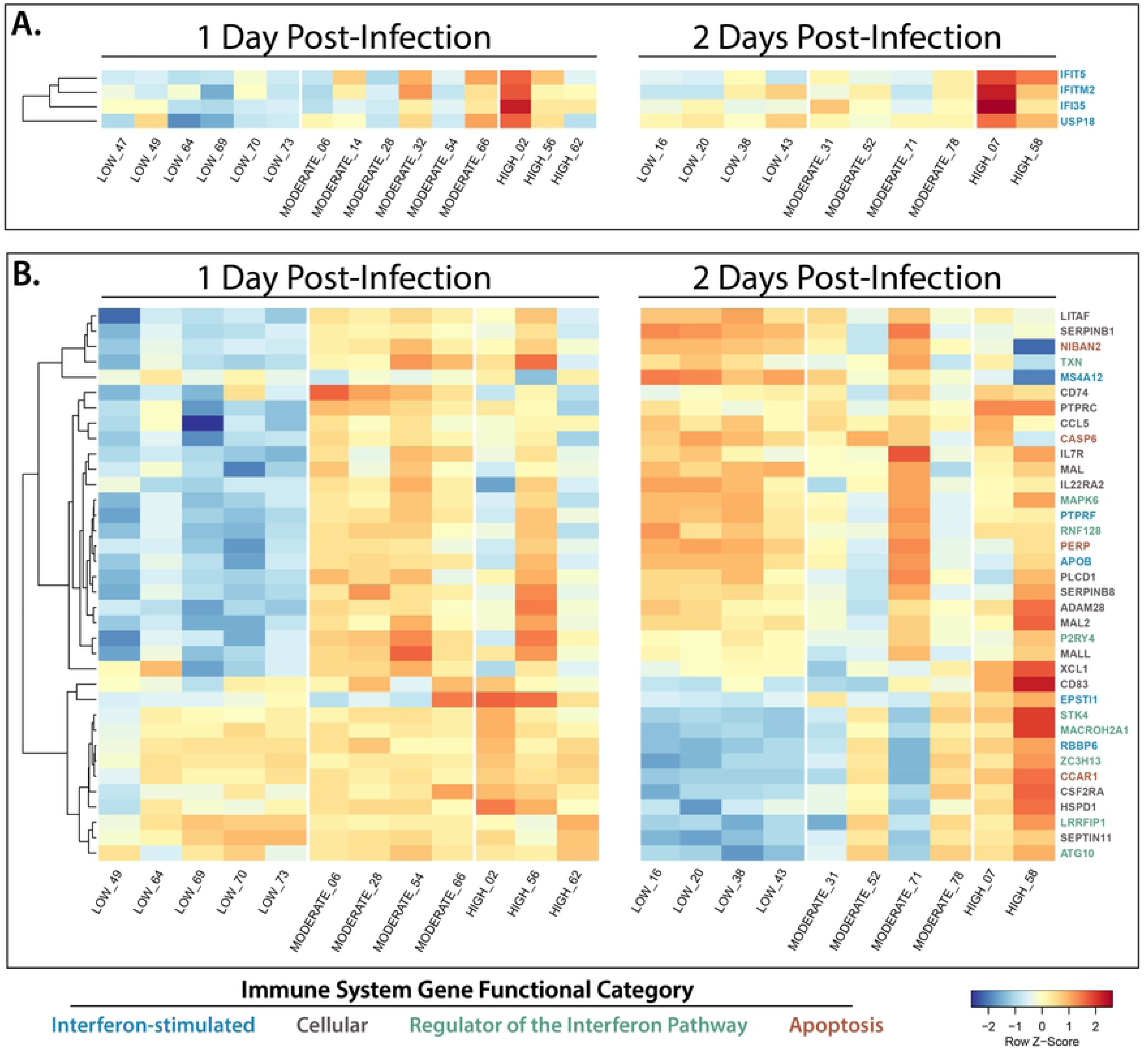
Immune genes differentially expressed in the bursa (A) and ileum (B) early in the infection. Individuals and their shed status group are on the x-axis and gene names are on the y-axis. Expression of genes is designated by color corresponding to the row Z-score (distance from the mean). Colors of gene names corresponds to their immune functional category.

**Table 2.**
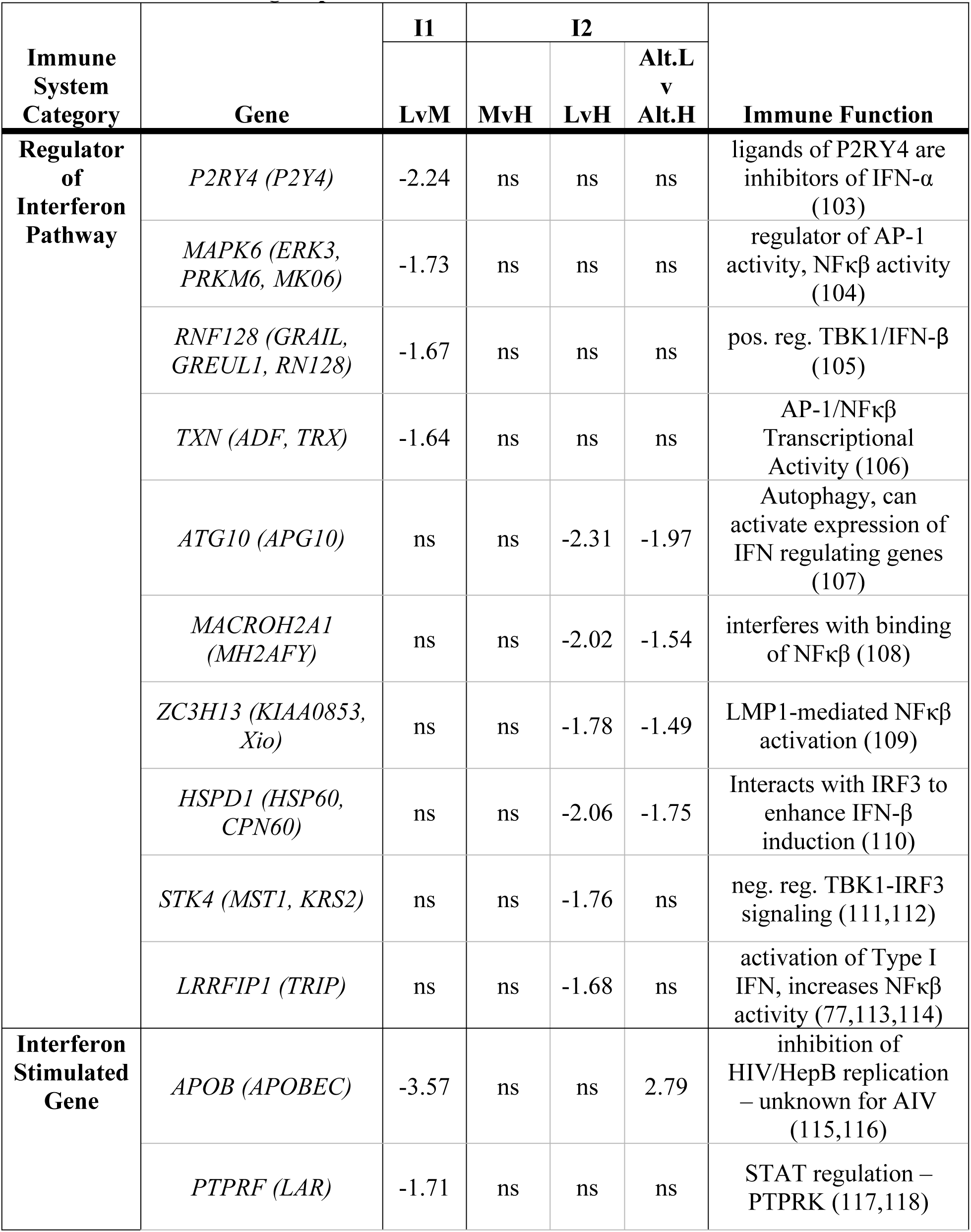
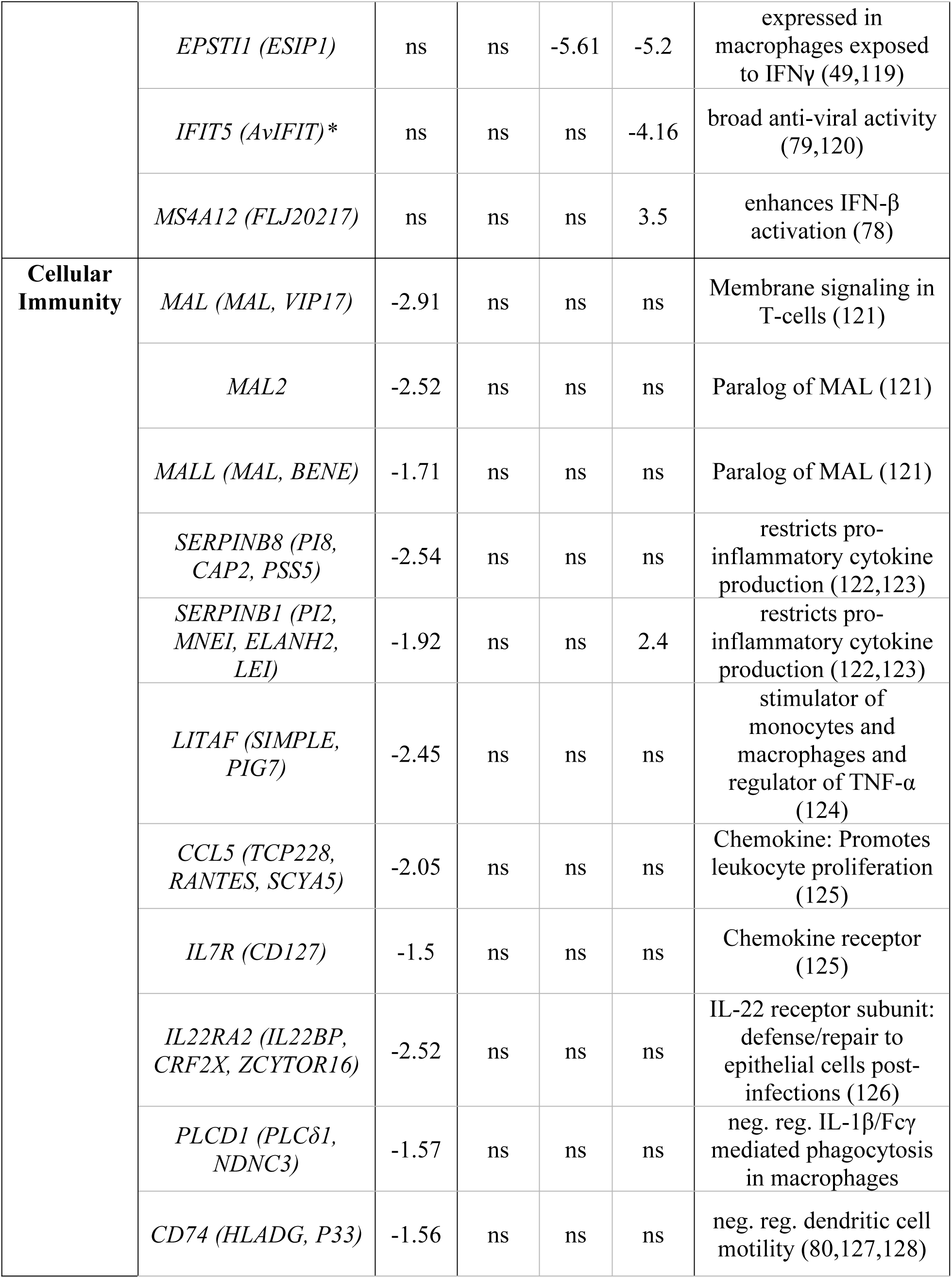
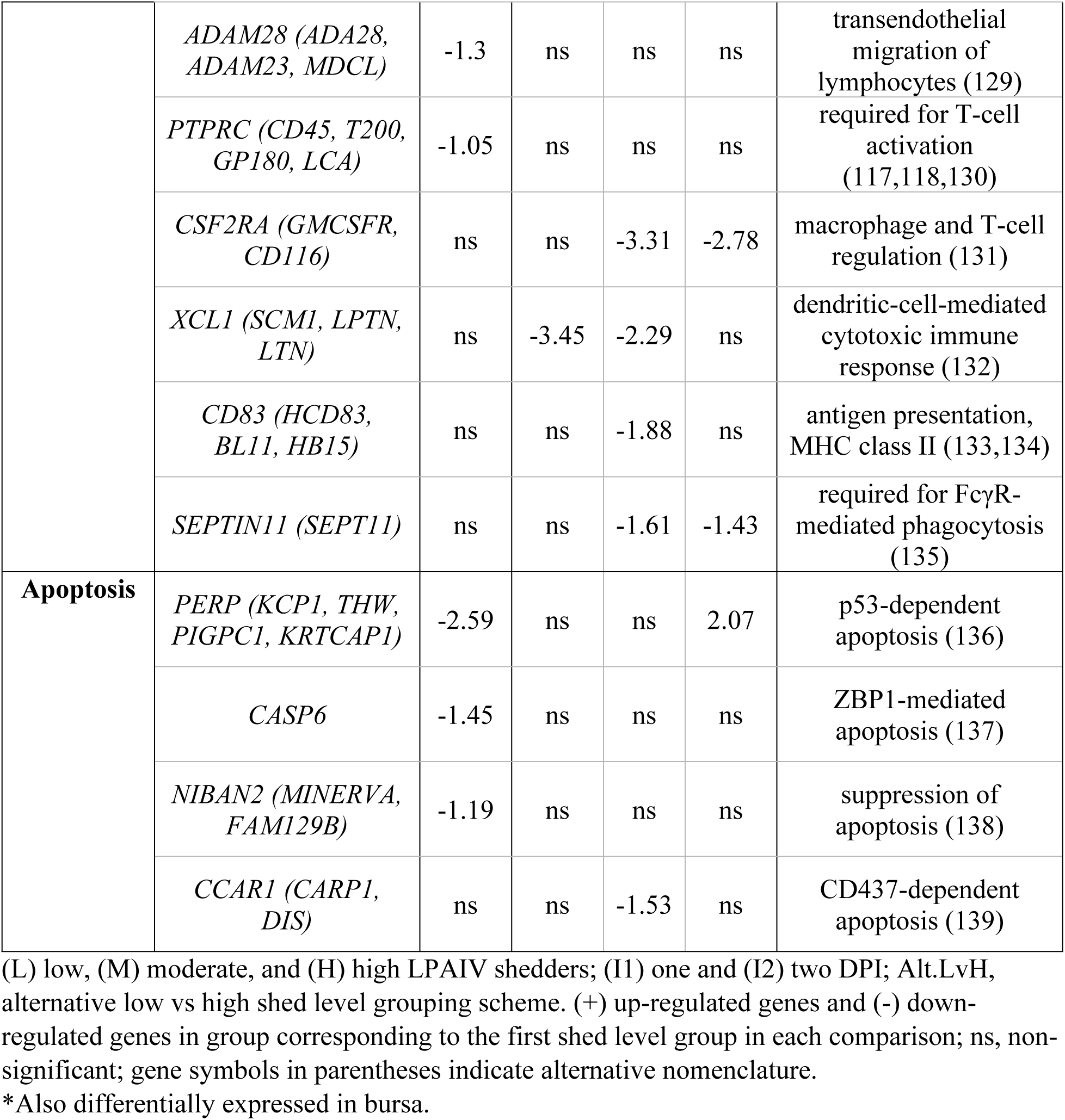
Log(fold change) differences for immune genes differentially expressed in the ileum between shed level groups.

**Table 3.** Log(fold change) differences for immune genes differentially expressed in the bursa between shed level groups.

| Immune System Category | Gene | I2 - LvH | Immune Function |
| --- | --- | --- | --- |
| **Interferon Stimulated Gene** | *IFIT5 (AvIFIT)\** | -6.57 | Broad anti-viral activity (120) |
|  | *IFITM2 (DSPA2c)* | -5.55 | Inhibition of viral replication by multiple mechanisms (79,140) |
|  | *USP18 (UBP18, ISG43)* | -3.56 | Neg. reg. type I IFN (141) |
|  | *IFI35 (IFP35)* | -3.36 | Neg. reg. RIG-I (142) |

(L) low and (H) high LPAIV shedders; (I2) two DPI; (-) down-regulated genes in group corresponding to the first shed level group in each comparison; gene symbols in parentheses indicate alternative nomenclature.
\*Also differentially expressed in ileum.

The significance of up-regulated immune genes in higher shedding birds early in the infection could indicate that mallards with higher viral shedding elicit stronger innate immune responses than low viral shedders. ISGs up-regulated in high shedding mallards, such as *IFIT5*, *EPSTI1*, *IFITM2*, *USP18*, and *IFI35*, were also up-regulated in AIV-infected poultry of previous studies when compared to uninfected poultry (41,49,143). These ISGs are specifically important for antiviral defense because they participate in various protective mechanisms that inhibit viral replication and regulate the interferon pathway (Table 2 and Table 3). ISG production is dependent upon a series of biochemical cascades beginning with PRRs recognizing intracellular virus which triggers the transcription of type I and II interferons (31,32,48), thus it is likely that intracellular virus is responsible for the interferon response. Although shed level groups were not characterized using intracellular virus titers, cloacal virus titers are known to be positively correlated with virus titers in ileum and bursa tissue (22); therefore, it is likely that high shedders also have high viral replication in these tissues. We suggest that beyond being activated by a viral infection, the magnitude of the innate immune response is dependent on high viral loads.

Due to the antiviral activity of ISGs, it could be hypothesized that high viral shedders with high ISG expression early in the infection may be able to clear the virus faster than low shedders. If so, then we would predict that high shedding birds early in the infection would have lower virus titers later in the infection. Given that tissue samples for gene expression were obtained when the bird was sacrificed, we were unable to assess their future viral shedding; however, for the birds in groups I5, I15, and I29, the virus titers early in the infection were positively related to virus shed later in the infection (S4 Table), which is consistent with other experimental infection studies (15). Previous findings also suggest that the up-regulation of interferon genes early in the infection does not lead to lower viral shedding (28, 144). Cornelissen et. al (28) did not find associations between interferon pathway genes and AIV clearance, and Helin et. al. (144) found that mallards continued to shed virus after observed up-regulation of interferon genes. For future studies, we recommend evaluating non-lethal samples for gene expression such as whole blood (145) or peripheral blood mononuclear cells (146) to gain further insight into the relationship between the immune response and viral shedding over time in individuals.

Interferon regulating genes associated with the production of ISGs, such as *DDX58*, *MDA-5*, *NFκβ*, *AP-1*, *IRF7*, *IFN-α*, *IFN-β*, and *IFN-λ,* that were previously differentially expressed between AIV-infected and uninfected poultry (28,39,41,43), were not differentially expressed in our study. We did however observe the up-regulation of *P2RY4*, *MAPK6*, *RNF128*, *ATG10*, *MACROH2A1*, *ZC3H13*, *STK4*, and *LRRFIP*, which all participate in regulating the interferon pathway (Table 2); therefore, the expression of genes involved in regulating the production of *DDX58*, *MDA-5*, *NFκβ*, *AP-1*, *IRF7*, *IFN-α*, *IFN-β*, and *IFN-λ* could impact the downstream expression of ISGs. These findings suggest that we may have identified key regulating components of the interferon pathway in LPAIV-infected mallards that are associated with viral shedding variation.

DEGs that are, or are related to, host cell factors (paralogs, orthologs) that promote viral replication were also found between shed level groups early in the infection (Table 4). Thirteen genes involved in promoting viral cell entry, including two sialyltransferases (*ST3GAL1* and *ST3GAL5*), were down-regulated in group I1 low shedders. These results are expected considering that *ST3GAL1* and *ST3GAL5* are enzymes responsible for the production of SAα2,3Gal (88), the viral binding receptor preferred by LPAIV to gain cell entry for replication (46). We previously detected a positive relationship between LPAIV shedding and SAα2,3Gal in ileum enterocytes of mallards (22), indicating that birds with fewer SAα2,3Gal had lower viral shedding quantities. Additionally, 11 genes that promote viral transcription, translation, and intracellular transport were up-regulated in group I2 high shedders. If viral cell entry is the key factor determining viral shedding on one DPI, it would make sense that genes promoting viral replication would be up-regulated in high shedding birds on two DPI. The observed differential gene expression of host cell factors between LPAIV shed level groups could explain why there is an observed one to two day latent period for AIV detection in poultry (147). Birds with fewer SAα2,3Gal may take longer to accumulate enough intracellular virus for replication, thus resulting in a delayed immune response. These findings suggest a connection between gene expression of host cell factors and immune genes in the ileum.

**Table 4.**
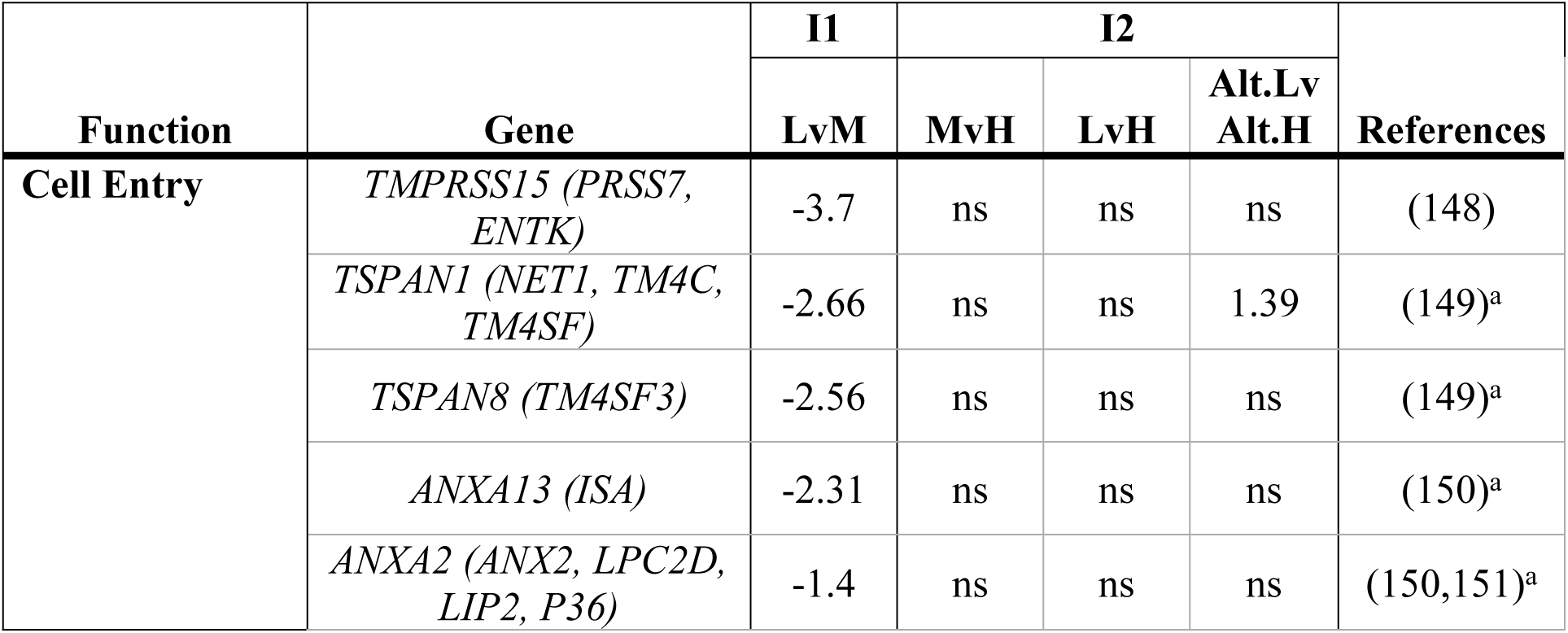
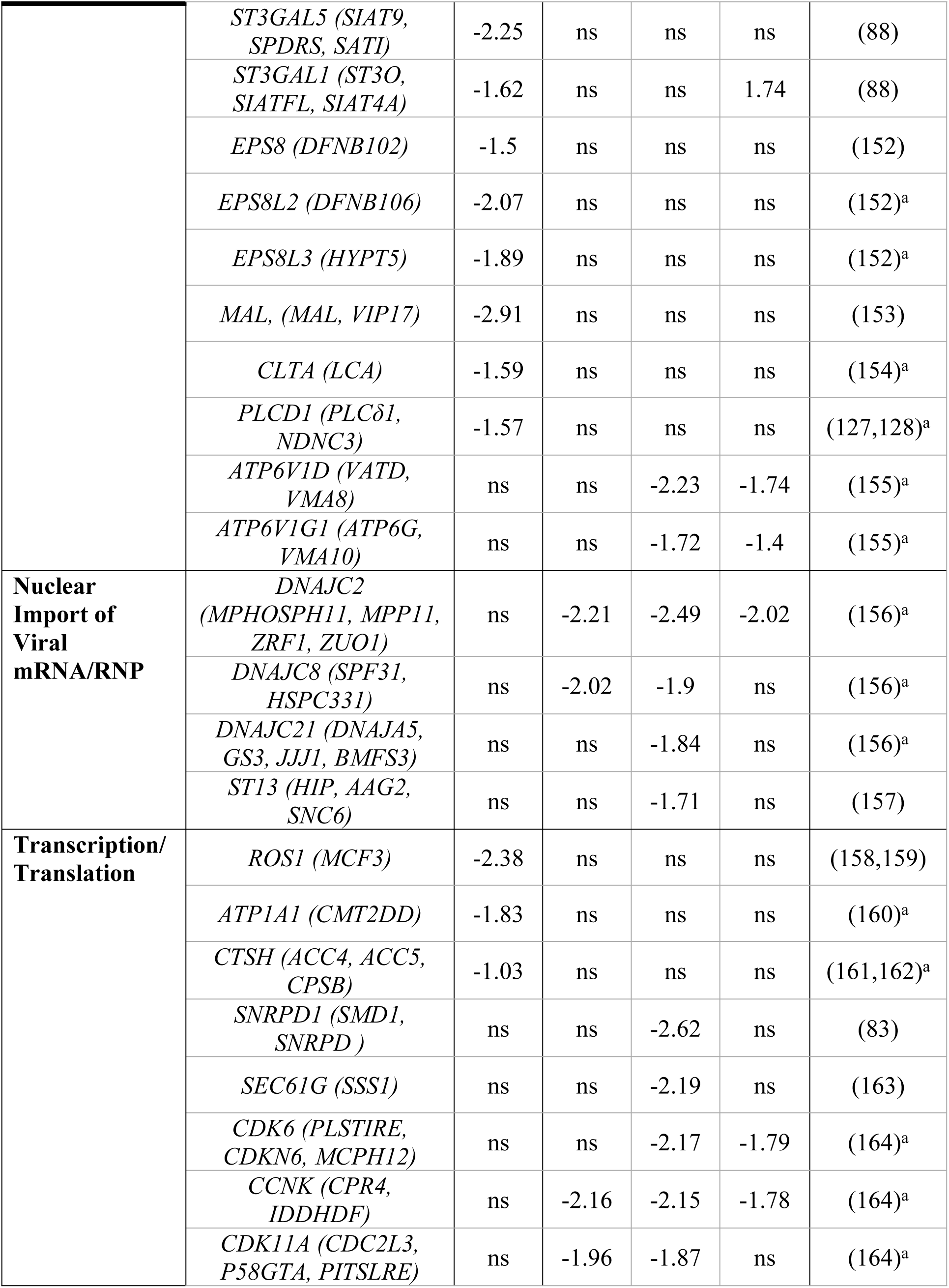
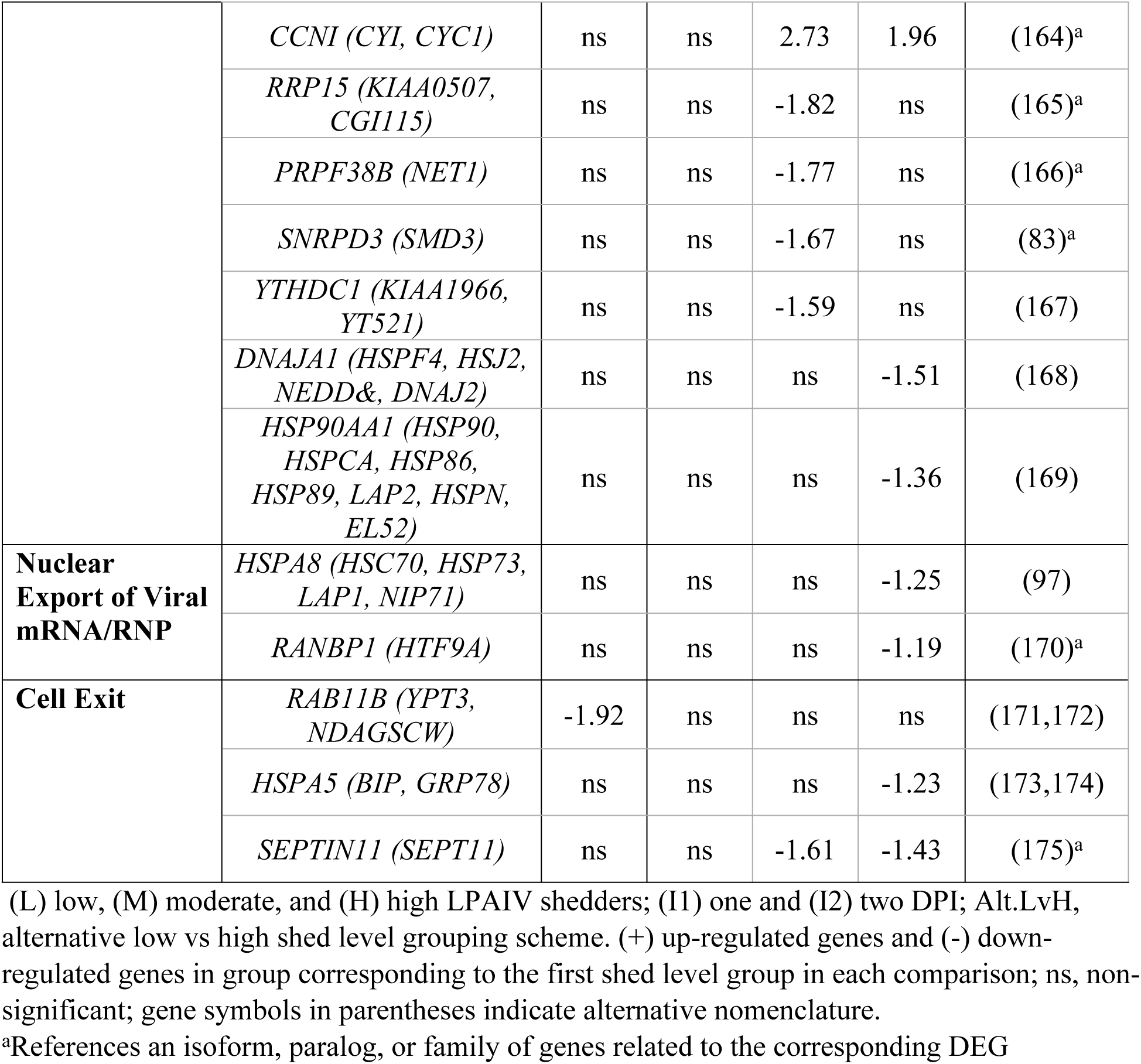
Log(fold change) differences for host cell factor genes that promote viral replication differentially expressed between shed level groups.

### Differential Gene Expression Between Virus Shed Level Groups During Late-Stage Infection

Contrary to our *apriori* predictions, differential expression was not observed between shed level groups during late-stage infection (group I5) at the gene or transcript level in the bursa or ileum (Section 7 in S2 Document). As stated above, genes associated with the adaptive immune response, including T-cell regulation and MHC class II expression were differentially expressed in the ileum early in the infection (Table 2). Our results suggest that immune cell mechanisms that lead to adaptive immunity, as it may relate to viral shedding, occurs earlier in the infection than we had expected.

### Candidate Gene Analysis

We assessed 268 candidate genes (Table B in S3 File), and 433 candidate transcripts (Table C in S3 File) for associations with viral shedding. Of those, 11 genes and seven transcripts in the bursa and thirteen genes in the ileum (transcripts were not assessed in ileum) had significant positive linear relationships with cloacal virus titers (S5 Document). No genes or transcripts had negative linear relationships. All the candidate genes with significant relationships to cloacal virus titers were related to the host immune system (Fig 9). Conversely, none of the sialyltransferase candidate genes assessed were related to cloacal virus titers. Expression levels of 11 out of the 15 genes were significantly associated with virus titers early in the infection (one and two DPI), but not late in the infection (five DPI).

**Fig 9.**
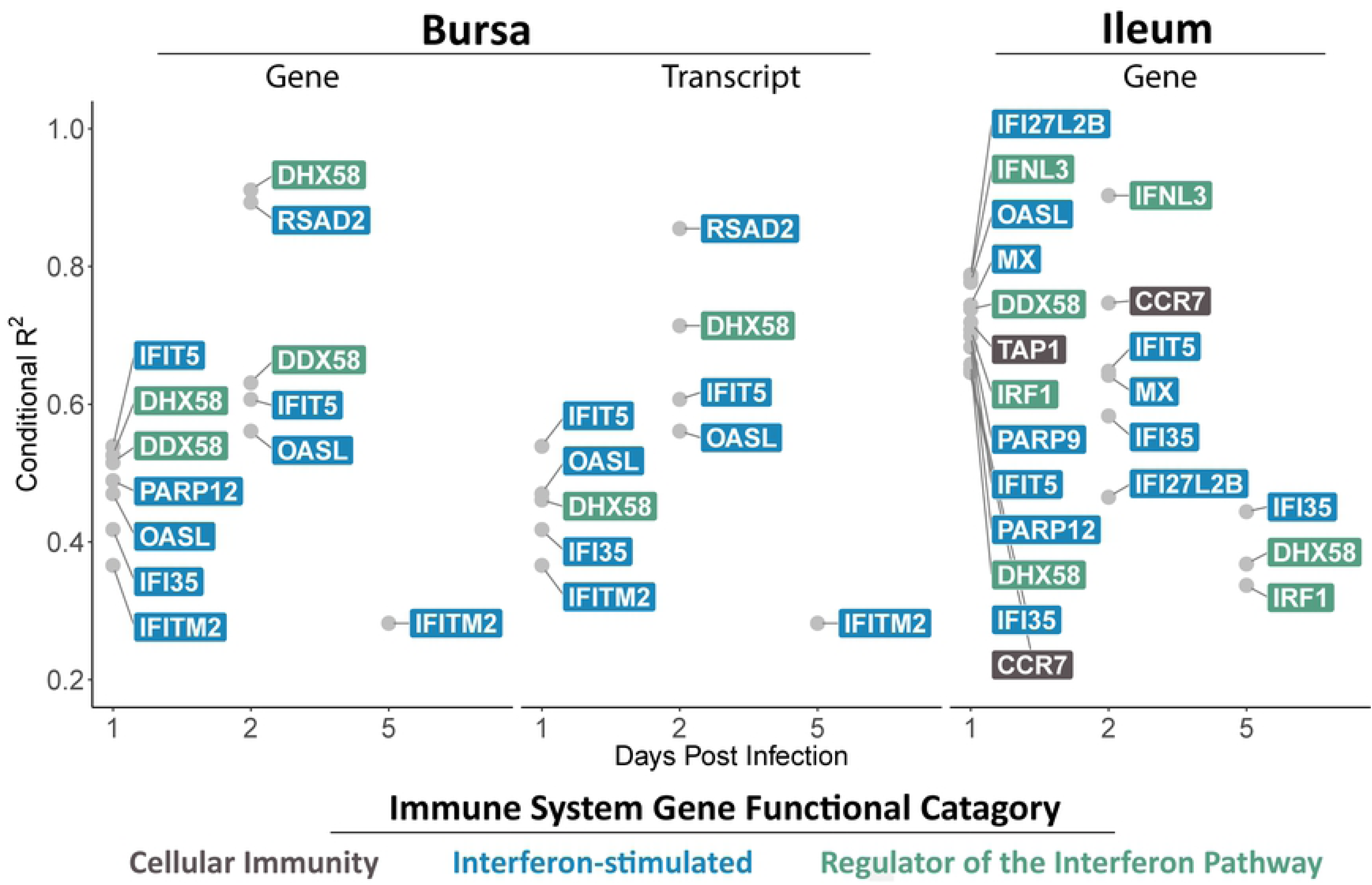
Conditional R^2^ values for candidate genes with significant (p <0.05) linear relationships to cloacal swab virus titers. Expression of all genes and transcripts shown are positively related to cloacal virus titers. Candidate genes in the bursa were evaluated at the gene and transcript level. Candidate genes in the ileum were only evaluated at the gene level. Colors of gene names corresponds to their immune gene functional category.

Interferon pathway genes up-regulated in previous LPAIV-infected ducks compared to uninfected ducks, such as PRRs *DDX58* and *DHX58,* and ISGs *IFIT5*, *IFITM2*, *OASL*, *RSAD2*, *MX*, and *PARP12* (40,42,85,87,176), were among the immune genes with positive relationships to viral shedding. Cornelissen et. al. (28) suggests that PRR gene expression is elevated in response to higher viral RNA levels, and Smith et. al. (41) suggests that *IFITM2* is strongly up-regulated in response to HPAIV. Collectively, these studies and our findings further suggest that viral shedding may be affecting the expression of interferon pathway genes.

Variation in viral shedding explained more variation in gene expression in the ileum than the bursa at one DPI. Similarly, our results are consistent with a previous study that found gene expression of PRRs was higher in the ileum compared to the bursa on one DPI (28). We also observed more immune DEGs in the ileum compared to the bursa between shed level groups (Fig 8, Table 2). Given that virus titers in the bursa and ileum are both positively related to cloacal swab virus titers with similar R^2^ values of 0.43 and 0.41 respectively (22), less viral replication in the bursa does not explain the closer relationship between viral shedding and immune gene expression in the ileum. Instead, we suggest the relationship between virus titers and gene expression in the ileum is explained by SAα2,3Gal in the ileum tissue, which we previously found to be positively associated with viral shedding (22). The down-regulation of sialyltransferase genes (*ST3GAL1* and *ST3GAL5*) in the ileum of low shedders in group I1 (Table 3) also supports this hypothesis. With fewer receptors (SAα2,3Gal) for the virus in the ileum, less virus is bound and replicated; therefore, the low viral shedding may induce a weaker innate immune response. Together, these findings may suggest that the ileum is more involved than the bursa in the relationship between the immune response and viral shedding variation.

### Conclusions and future directions

Intraspecific variation in pathogen shedding can alter the magnitude and duration of infectious disease outbreaks; hence, understanding the host intrinsic factors associated with variation in pathogen shedding is key for developing disease management plans. Here, we evaluated the variation in tissue-specific gene expression as it relates to infection status, DPI, and pathogen shedding in LPAIV-infected mallards to obtain a better understanding of the molecular mechanisms underlying intraspecific variation in pathogen shedding. Cloacal viral shedding was measured because it represents transmissible virus and sample collection can be repeated over time (58), thus providing a direct link to questions of transmission dynamics. Gene expression was evaluated in the ileum and bursa because these tissues are sites of high LPAIV replication in mallards (44,45,47). Our findings not only provided evidence that gene expression in wild mallards is more associated with viral shedding than by infection status alone, but also provided insight into the host cell machinery utilized by the virus to successfully replicate. These data improve the current understanding of avian host genes involved in LPAIV-infections and the molecular mechanisms associated with viral shedding magnitude.

Collectively, we have identified specific genes and biological processes in the ileum and bursa of LPAIV-infected mallards that are associated with viral shedding heterogeneity; however, this study alone does not confirm an inherent genetic difference between high viral shedders and low viral shedders. Additionally, while we show a relationship between viral shedding and gene expression, we were unable to determine whether viral shedding influenced gene expression or vice versa, which also makes it difficult to determine an inherent cause and effect. Single nucleotide polymorphisms (SNPs) in relevant genes could be accounting for the differential gene expression observed; however, the gene mapping methods we used to detect gene expression did not account for SNP differences (66). A potential next research step would be to identify SNPs in observed DEGs to determine if a specific gene SNP is associated with intraspecific viral shedding variation (177). For example, a specific allele identified in cattle has been associated with lower proviral loads of bovine leukemia virus (178). When BLV-infected cattle selected for this specific low proviral allele were introduced to uninfected dairy herds, viral transmission stopped (26). We recommend that the genes identified in our current study, such as interferon pathway genes *DDX58*, *LGP2*, *IFIT5, IFITM2, IFI35, FOSB* (Fig 10), viral replication promoting genes *HSPA8*, *ST3GAL1, ST3GAL5,* and other genes described in Tables 2, 3, and 4, be examined further by identifying SNPs, utilizing gene-knockout studies, and conducting site-directed mutagenesis to understand the potential for inherent genetic differences (e.g., gene mutations) and associated molecular mechanisms that may contribute to viral shedding variation.

**Fig 10.**
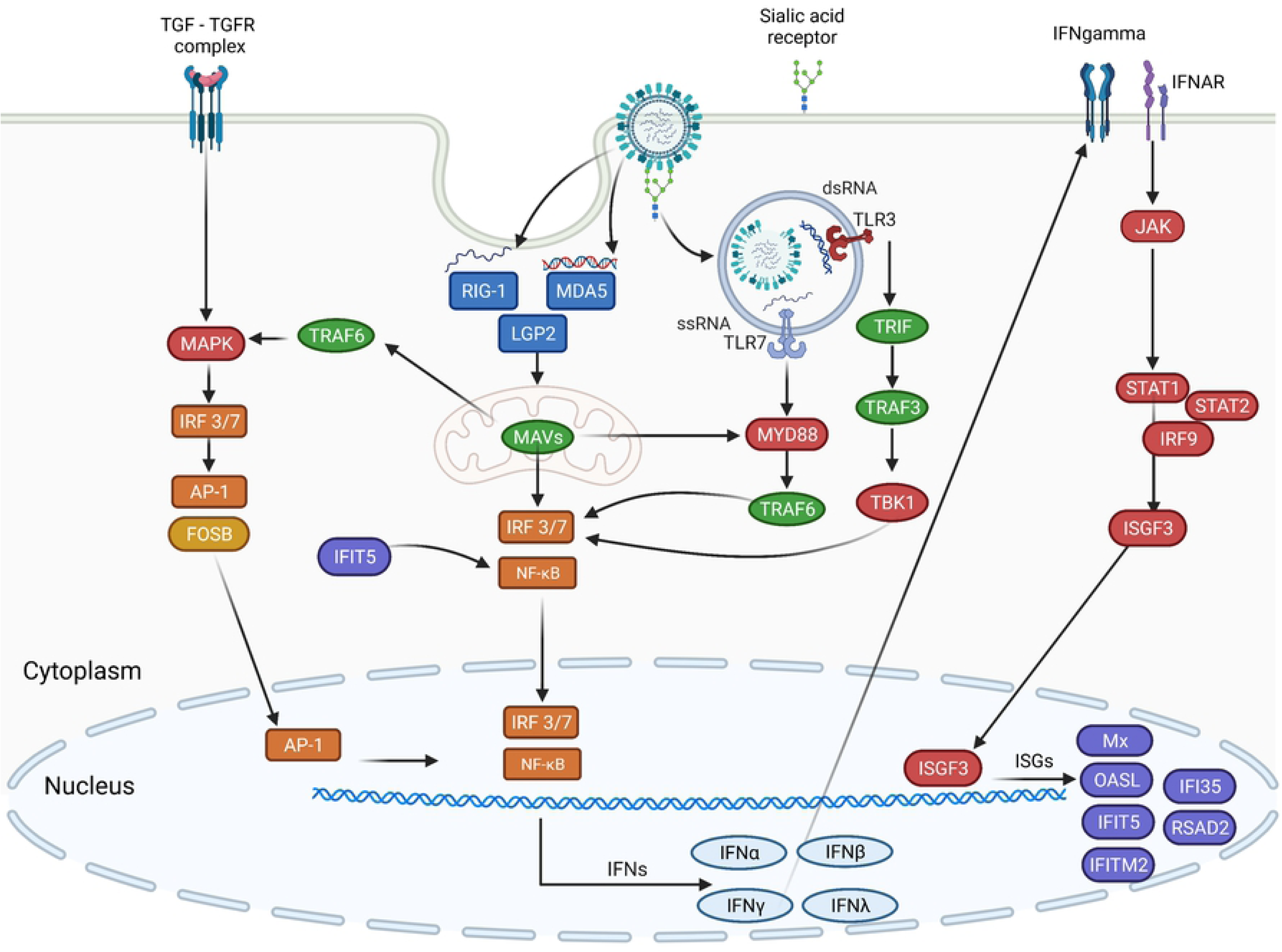
Intracellular innate immune response to avian influenza. Antiviral interferon stimulated genes (ISGs, purple) are produced through a cascade of intracellular signaling processes (signaling molecules are in red) known as the interferon pathway. Pattern recognition receptors (dark blue) and toll-like receptors (TLRs) stimulate adaptor molecules (green) for the production of transcription factors (orange). Transcription factors relocate to the nucleus to trigger the production of interferons (IFN, light blue). Interferons transport outside the nucleus to initiate the JAK-STAT signaling pathway responsible for the production of the ISGF3 complex, which relocates to the nucleus to stimulate the production of ISGs. ISGs have various antiviral effects targeting viral replication or directly acting on the interferon pathway. Created with BioRender.com.

The data acquired in this study should also be used to determine how the intrinsic host factors associated with viral shedding contribute to pathogen transmission. We know that identifying and removing super-shedders is a method of controlling outbreaks as evidenced by modeling the super-shedders of *E.coli* O157-infected cattle (179). If the individuals with the propensity to be the super-shedders can be determined prior to pathogen exposure, new strategies for outbreak prevention could be realized. This future research depends on applying the knowledge gained from exploratory studies like this one, as well as performing similar studies in multiple host-pathogen systems, to obtain a better understanding of how individual host variation contributes to the complexity of disease transmission dynamics. With more knowledge gained, the more tools wildlife managers, veterinarians, and public health officials will have to better predict and control future outbreaks.

## Supporting information

**S1 Table. RNA integrity number (RIN) and sequencing pool for mRNA samples isolated from ileum and bursa tissue.**

**S2 Document. Complete differential gene expression results for all group comparisons tested in ileum and bursa tissue.**

**S3 File. Excel spreadsheet with candidate gene search terms (Table A), candidate genes (Table B), and candidate transcripts (Table C).**

**S4 Table. Correlation matrix with Pearson’s R correlation coefficients for cloacal swab virus titers on 1-5 days post infection.**

**S5 Document. Complete candidate gene analysis results showing positive linear relationships between candidate genes in the ileum and candidate genes/transcripts in the bursa.**

## Acknowledgements

We thank F. Rohwer and M. Buxton from Delta Waterfowl for assistance in egg collection; J.P. Steibel for assistance with RNAseq experimental design; staff at the MSU genomics center for library sequencing, RNAseq, and assistance with RT-PCR; staff at the UMN genomics center for library sequencing and RNAseq; D. Incorvaia and E. Nelson, who were NSF funded undergraduate interns (REU); and E. Hergert, C. Sklarczyk, M. Yang, T. McCord, M. Phillips, O. Lefere, W. Decker, V. Olszewski, and V. Colberg who were undergraduate volunteers who assisted with the collection of eggs, care of birds, sample collection, and laboratory sample processing.

